# Genomic and phenotype-based framework for rational design of phage cocktails against *Escherichia coli* isolates from urinary tract infections

**DOI:** 10.64898/2026.09.24.754182

**Authors:** Paul Ugalde Silva, Ally Champoux, Annie Chénard, Serena Tuytschaevers, Élodie Jolicoeur, Nathalie Rivard, Louis-Charles Fortier

## Abstract

Urinary tract infections (UTIs) are frequently caused by *Escherichia coli* and are increasingly associated with antimicrobial resistance, highlighting the need for alternative therapies. Although phage therapy is a promising approach, the rational selection of phages and phage cocktails remains challenging, particularly when detailed genomic information on the infecting strain is unavailable. In this study, we isolated 18 phages from environmental samples collected in Sherbrooke, Quebec, Canada. Genome sequencing identified 15 pure lytic phages representing diverse taxonomic groups and revealed prophage-derived contaminants in three isolates, underscoring the importance of genomic validation during phage discovery. Host range was determined using efficiency-of-plating (EOP) assays against UTI-associated *E. coli* isolates. Based on host-range overlap and cumulative EOP values, we developed a phenotype-based framework for phage cocktail design and classified combinations as redundant or complementary. Complementary cocktails composed of phages with distinct host ranges and high infection effectiveness consistently outperformed redundant combinations in time-kill assays, demonstrating that host-range diversity and lytic effectiveness are independent properties that should be considered separately during cocktail design. To evaluate cocktail performance under physiologically relevant conditions, the most effective combinations were tested in artificial urine. Both cocktails exhibited reduced activity compared with standard laboratory media, indicating that environmental conditions strongly influence phage-host interactions. Together, our results demonstrate that host-range overlap and cumulative infection effectiveness provide a practical framework for identifying effective phage combinations in the absence of detailed bacterial genomic information and may facilitate the rational development of phage therapies for UTIs.

## Importance

The increasing association between urinary tract infections (UTIs) and antimicrobial resistance highlight the need for alternative therapeutic approaches. In this study, we isolated and characterized 15 novel lytic phages targeting *Escherichia coli* isolates from UTI patients. The novelty of this work lies in the integration of comprehensive phage genomic characterization with the development of a phenotype-based framework for phage cocktail design. While genome sequencing was used to validate phage identity, determine phage isolates purity, and characterize genomic diversity, cocktail selection was based exclusively on experimentally derived infection profiles, including host-range overlap and cumulative infection effectiveness. By using this framework, we demonstrated that phage cocktails composed of functionally complementary phages outperform redundant combinations, despite the absence of detailed genomic information on the target bacteria. The results of this study contribute new phage resources and describe a practical framework that may facilitate the rational development of phage cocktails for the treatment of UTIs.

## Introduction

Urinary tract infections (UTIs) constitute one of the most prevalent diseases worldwide, representing a substantial clinical and public health burden across both community and healthcare environments. UTIs can occur in individuals of any age or sex. However, their prevalence in women is significantly higher than in men (He et al., 2025; Yang et al., 2022). In 2019, an estimated 405 million cases were reported worldwide, including approximately 236,000 deaths, representing a 2.4-fold increase in mortality compared with 1990 (Yang et al., 2022).

UTIs are classified as either uncomplicated or complicated, depending on the patient’s underlying health status. Uncomplicated UTIs occur in individuals who are otherwise relatively healthy, whereas complicated UTIs arise in patients with additional conditions such as functional abnormalities of the urinary tract, other comorbid illnesses, immunocompromised states, or long-term catheterization (Nielubowicz & Mobley, 2010) . These infections are caused by both Gram-negative and Gram-positive bacteria, as well as by certain fungi species. Some pathogenic strains of *Klebsiella pneumoniae*, *Staphylococcus saprophyticus*, *Enterococcus faecalis*, group B *Streptococcus* (GBS), *Proteus mirabilis*, *Pseudomonas aeruginosa*, *Staphylococcus aureus* and *Candida* spp. are prevalent in uncomplicated UTIs. While *Enterococcus* spp., *K. pneumoniae*, *Candida* spp., *S. aureus*, *P. mirabilis*, *P. aeruginosa* and GBS are recognized causative agents of complicated UTIs. However, the most frequent pathogen causing both uncomplicated and complicated UTIs is uropathogenic *Escherichia coli* (UPEC) (Flores-Mireles et al., 2015) .

At present, antibiotic treatments stand as the prevailing and recommended therapeutic option for UTIs. However, the escalating rate of antibiotic resistance, coupled with a high recurrence rate, constitutes challenges to their effective treatment (Flores-Mireles et al., 2015). The emergence of antibiotic-resistant UPECs, as well as the spread of MDR (multi-drug resistant) UPEC strains, is a current clinical problem, especially for women with recurrent UTIs. The indiscriminate use of broad-spectrum antibiotics, especially in developing countries, has increased the frequency of cases caused by MDR strains of UPEC. In developing countries, resistance of UPEC to fluoroquinolones such as ciprofloxacin, commonly used for the treatment of UTIs, ranges from approximately 55% to 86%, while in developed countries it ranges from 5 to 39%. Resistance to trimethoprim–sulfamethoxazole is similarly high, at around 54% to 82% in developing countries, compared with approximately 15% to 39% in developed countries (Kot, 2019) . Combined with the decreasing numbers of new antibiotics discovered, this causes problems for treatment of UTIs, therefore research is turning toward alternative antimicrobial agents.

Bacteriophages (or phages), viruses that infect bacteria, have been used as therapeutic agents to combat bacterial infections for over a century in Eastern Europe and Russia but their use in Western countries was supplanted by antibiotics. However, with the global rise in antimicrobial resistance, there is now a renewed interest for phage therapy in Western countries (Gordillo Altamirano & Barr, 2019) .

Phages play an important role in the organization of microbial niches, exerting distinct influences depending on their infective cycle (Hobbs & Abedon, 2016) . All phages initiate infection by binding to specific receptors at the bacterial surface, followed by the injection of their genomes into the cytoplasm. After this step, their infective strategies fall into two broad categories: temperate phages, which alternate between dormant and productive states, and virulent phages, which follow a strictly lytic, productive cycle, leading to the death of the host. Due to their characteristics, lytic phages are an interesting option for phage therapy (Davies et al., 2016).

Phage therapy offers several advantages over antibiotics, including high specificity, self-replication, auto-dosing, low toxicity, and adaptability (Loc-Carrillo & Abedon, 2011; Subedi et al., 2025). As natural predators, phages selectively target limited bacterial genera, species, and even specific strains, thereby minimizing disruption of the surrounding microbiota in contrast to antibiotics (Emencheta et al., 2023) . Host specificity is primarily determined by the interaction between bacterial surface receptors and phage receptor-binding proteins (RBPs) (Klein-Sousa et al., 2025) .

Because antibiotics and phages employ distinct mechanisms to kill bacteria, antibiotic resistance may not confer cross-resistance to phages. Consequently, phages are also capable of targeting MDR bacteria (Loc-Carrillo & Abedon, 2011) .

Currently, the compassionate use of therapeutic phages is being applied to a variety of clinical conditions, including UTIs. Previous and ongoing studies indicate that phage therapy for UTIs can effectively reduce microbial load and alleviate clinical symptoms without raising concerns about significant side effects. These outcomes have been observed using either single phages or phage cocktails (Al-Anany et al., 2023), as well as in efforts to combat biofilm formation on clinical devices (Pawłuszkiewicz et al., 2025) .

One problem with phage therapy is that it can be a double-edged sword. On one hand, phage specificity is an advantage for protecting the surrounding microbiota. But on the other hand, this specificity can be a problem when the target strain cannot be infected by the available phages. Accordingly, it is essential to establish a substantial phage bank to ensure a wide range of therapeutic options for this type of infection, whether using individual phages or phage combinations in cocktails. Furthermore, in clinical settings, genomic information of the infecting pathogen is often unavailable when a therapeutic cocktail must be selected. At the same time, RBP are highly diverse, and both their sequences and receptor specificities remain poorly characterized. Consequently, predicting phage host range based solely on genomic information remains challenging, limiting the development of fully rational strategies for phage cocktail design (Gaborieau et al., 2024; Klein-Sousa et al., 2025).

The discovery and characterization of new phages represent key steps toward streamlining the application of phage therapy in the treatment of UTIs. In this study, we isolated and characterized 18 novel phages from diverse environmental sources from Sherbrooke surroundings (Quebec, Canada), including rivers, streams, swamps, and wastewater treatment plants. By combining host-range determination, efficiency-of-plating measurements, time-kill assays, and testing in artificial urine, we established a phenotype-based framework for phage cocktail design based on host-range overlap and infection effectiveness. This approach enabled the identification of complementary and redundant phage combinations, the selection of highly effective cocktails against *E. coli* strains from UTI patients, and the evaluation of their performance under conditions that more closely mimic the urinary tract environment. We also demonstrate that rigorous phage genome sequencing can reveal significant host-derived prophage contaminations that should be taken into consideration during the development of phage therapeutics.

## Results

### Phage isolation and plaque morphology

The isolated phages, their corresponding host strains, and the environmental sources from which they were recovered are summarized in Table 1. Phage isolation and propagation were performed using three *E. coli* strains: UTI strains 1278 and 1274, and the commensal mouse strain 1566. The inclusion of a mouse strain aimed to increase the likelihood of isolating a broader diversity of phages. In total, 18 phages were initially recovered from sewage water, swamps, and rivers in Sherbrooke, Quebec, Canada. Of these, nine were propagated using strain 1278, eight using strain 1566, and one using strain 1274. Genome sequencing subsequently revealed that three isolates contained host-derived prophage contaminants and were therefore excluded from further analyses (Supp Table 2). The remaining 15 purified lytic phages constituted the final dataset used for genomic characterization and host-range studies. All pure isolated phages produced lysis zones and plaques in spot tests on their respective host strains. Plaques were typically small (∼1 mm) and well defined (Fig. 1).

**Figure 1.**
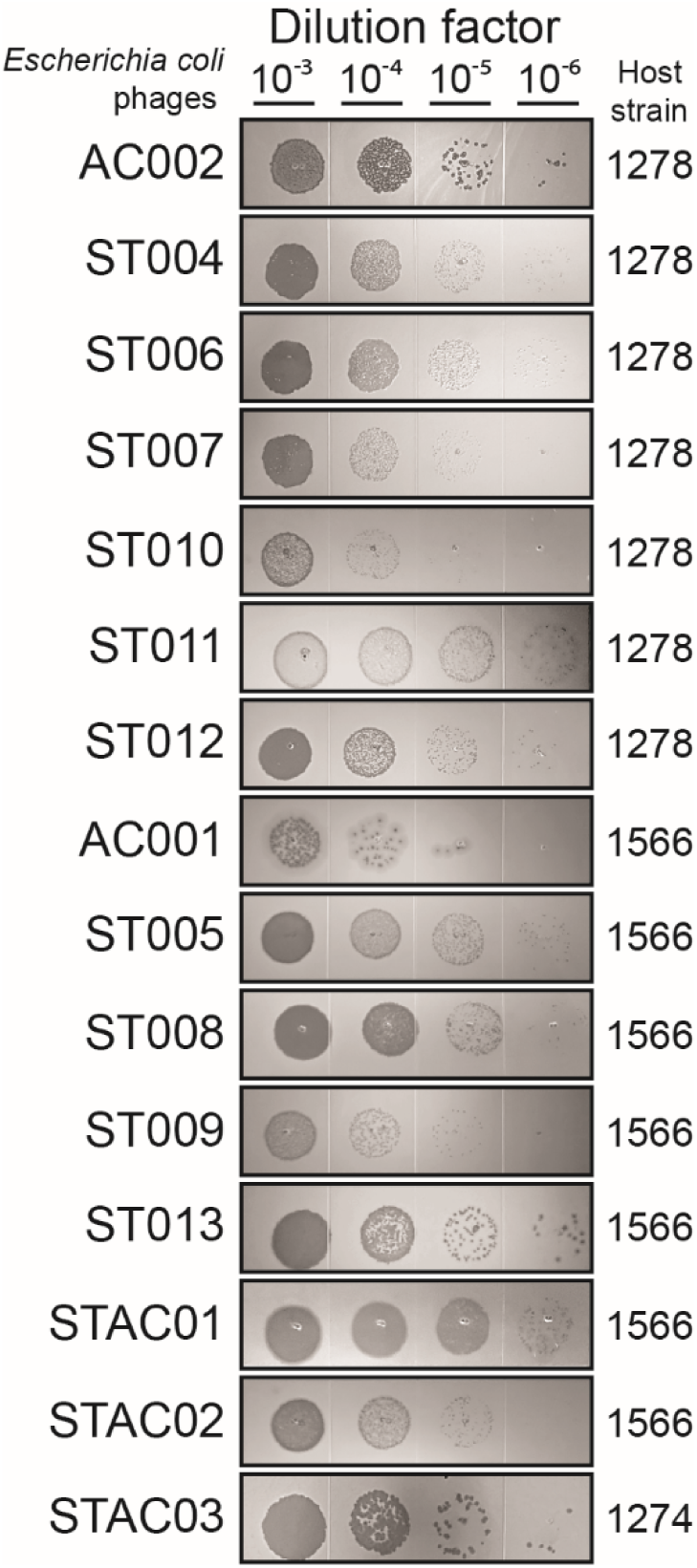
Spot test assay with UTI strains 1278 and 1274, and the commensal mouse strain 1566. 10-fold dilutions of each isolated *Escherichia coli* phage were spotted on top of bacterial lawn of the corresponding host strain. Clear zones indicate bacteria lysis.

**Table 1.** Environmental phages isolated from different locations in Sherbrooke, Quebec, Canada.

| *Escherichia coli* phage | Origin | Host strain |
| --- | --- | --- |
| AC002 | Wastewater treatment plant | 1278 |
| ST004 | Wastewater treatment plant |  |
| ST006 | Wastewater treatment plant |  |
| ST007 | Wastewater treatment plant |  |
| ST010 | Swamp water |  |
| ST011 | River near a dam |  |
| ST012 | Wastewater treatment plant |  |
| AC001 | Wastewater treatment plant | 1566<br>(mouse) |
| ST005 | Wastewater treatment plant |  |
| ST008 | Vegetable fields |  |
| ST009 | Swamp water |  |
| ST013 | Contaminated stream |  |
| STAC01 | River under a bridge |  |
| STAC02 | Swamp water |  |
| STAC03 | Wastewater treatment plant | 1274 |
| Host-derived prophage contaminated samples |  |  |
| ST014 | Wastewater treatment plant | 1566 |
| AC003 | Wastewater treatment plant | 1278 |
| AC004 | Wastewater treatment plant | 1278 |

**Table 2.** Bioinformatic analysis of the isolated phages.

| *Escherichia coli* phage | GenBank accession | Genome size (bp) | CDS | Hypothetical proteins | Lysogeny-associated genes | Closest relative (INPHARED) | Lowest taxa (genus) | Identity (%) | Coverage (%) | GC content (%) |
| --- | --- | --- | --- | --- | --- | --- | --- | --- | --- | --- |
| AC002 | PZ938575 | 40,632 | 62 | 14 | 2 | *Escherichia* phage vB_EcoP_ZX5 | *Uetakevirus* | 99.14 | 81 | 48.98 |
| ST004 | PZ938564 | 150,491 | 309 | 120 | 0 | *Escherichia* phage ESCO13 | *Phapecoctavirus* | 99.05 | 95 | 39.12 |
| ST006 | PZ938566 | 152,873 | 314 | 120 | 0 | *Escherichia* phage ESCO13 | *Phapecoctavirus* | 98.61 | 95 | 39.12 |
| ST007 | PZ938567 | 151,093 | 308 | 123 | 0 | *Escherichia* phage ESCO13 | *Phapecoctavirus* | 97.93 | 95 | 39.12 |
| ST010 | PZ938570 | 150,485 | 306 | 114 | 0 | *Escherichia* phage PNJ1809-36 | *Phapecoctavirus* | 99.59 | 97 | 39.12 |
| ST011 | PZ938571 | 150,358 | 312 | 124 | 0 | *Escherichia* phage vB_EcoM-Ro121lw | *Phapecoctavirus* | 99.39 | 99 | 39.08 |
| ST012 | PZ938572 | 153,644 | 313 | 121 | 0 | *Escherichia* phage ESCO13 | *Phapecoctavirus* | 97.88 | 95 | 39.12 |
| AC001 | PZ938574 | 41,583 | 55 | 13 | 0 | *Serratia* phage vB_SmaP-Kaonashi | *Aghbyvirus* | 98.4 | 98 | 53.92 |
| ST005 | PZ938565 | 152,789 | 313 | 120 | 0 | *Escherichia* phage ESCO13 | *Phapecoctavirus* | 98.6 | 95 | 39.12 |
| ST008 | PZ938568 | 133,904 | 264 | 109 | 0 | *Escherichia* phage vB_EcoM_TH18 | *Avunavirus* | 96.12 | 93 | 39.95 |
| ST009 | PZ938569 | 150,185 | 312 | 126 | 0 | *Escherichia* phage vB_EcoM-Ro121lw | *Phapecoctavirus* | 98.75 | 98 | 39.08 |
| ST013 | PZ938573 | 151,210 | 308 | 121 | 0 | *Escherichia* phage ESCO13 | *Phapecoctavirus* | 97.93 | 95 | 39.12 |
| STAC01 | PZ938576 | 52,432 | 84 | 22 | 0 | *Salmonella* phage emhyr | *Rosemountvirus* | 95.66 | 99 | 45.89 |
| STAC02 | PZ938577 | 136,288 | 252 | 99 | 0 | *Escherichia* phage nomo | *Vequintavirus* | 98.35 | 99 | 43.65 |
| STAC03 | PZ938578 | 167,514 | 286 | 75 | 0 | *Escherichia* phage vB-Eco-KMB26 | *Tequatrovirus* | 99 | 98.96 | 35.33 |

### Genome sequencing and bioinformatic analysis of isolated phages

The complete genomes of the 18 isolated phages were sequenced using Oxford Nanopore Technology. Genome annotation was performed with Pharokka (version 1.8.2)) (Bouras et al., 2023) and Empathi (Bouras et al., 2026), and the lowest taxonomic rank (genus) and the closest relatives were identified by using Pharokka through the Mash algorithm with the INPHARED database (version 1.2) (Table 2 and Supp Table 2). Nine phages were identified as members of the genus *Phapecoctavirus*, displaying a relatively uniform genome size of approximately ∼150 kb with a GC content of ∼39%. The remained six phages belonged to the genera such as *Uetakevirus*, *Aghbyvirus*, *Avunavirus*, *Rosemountvirus*, *Vequintavirus*, and *Tequatrovirus*, and exhibited substantial variation in genome size and GC content. Among these, the *Uetakevirusi* member had the smaller genomes (40.632 kb with a 48.98% GC content), where as the *Tequatrovirus* member possessed the larger genome (167.514 kb with a 35.33% GC content).

The initial phage collection comprised 18 isolates. However, genome analysis revealed that three of the initially isolated phages corresponded to mixtures of target phages and contaminant elements, including prophages or episomal phages likely derived from the host strains and introduced during propagation. Most of these elements contained one or two lysogeny-associated genes, suggesting a temperate lifestyle. These three phages and their corresponding genomes were therefore excluded from further analyses (Supp. Table 2).

Annotation using Pharokka and Empathi identified approximately 310 coding domain sequences (CDSs) in phages ΦST005, ΦST006, ΦST012, ΦST011, ΦST013, ΦST007, ΦST009, ΦST010, and ΦST004, from which ∼240 were annotated as hypothetical proteins with unknown function. In contrast, 252 and 264 CDSs were identified in phages ΦSTAC02 and ΦST008, respectably, including 181 and 172 hypothetical proteins. Phage ΦSTAC03 contained 286 CDSs, among which 148 were annotated as hypothetical proteins. For phages ΦSTAC01, ΦAC002, and ΦAC001, 84, 62, and 55 CDSs were identified, respectively, with 43, 39, and 29 corresponding to hypothetical proteins.

Genome alignment of the 15 phages using GenALLYgn (Version 1.0) revealed that nine phages belonging to the genus *Phapecoctavirus* clustered together and shared a substantial number of genes. Phages ΦSTAC02 and ΦST008 shared only a limited number of genes with the *Phapecoctavirus* members. In contrast with ΦSTAC03, ΦSTAC01, ΦAC002, and ΦAC001, did not shared any detectable genes with these groups, indicating that they are genetically more distant (Fig. 2A). Intergenomic similarity score (ISS) calculated by VIRIDIC further supported these relationships. The *Phapecoctavirus* phages exhibited the highest ISS values, ranging from 92% to 99.5%. Phages ΦSTAC02 and ΦST008 showed an ISS value of 14.8%, consistent with the limited number of shared genes observed in the genome alignment. In contrast, the four genetically distant phages displayed ISS values ranging from 0% to 3.8%, confirming their high level of divergence from the rest of the phage collection (Fig. 2B).

**Figure 2.**
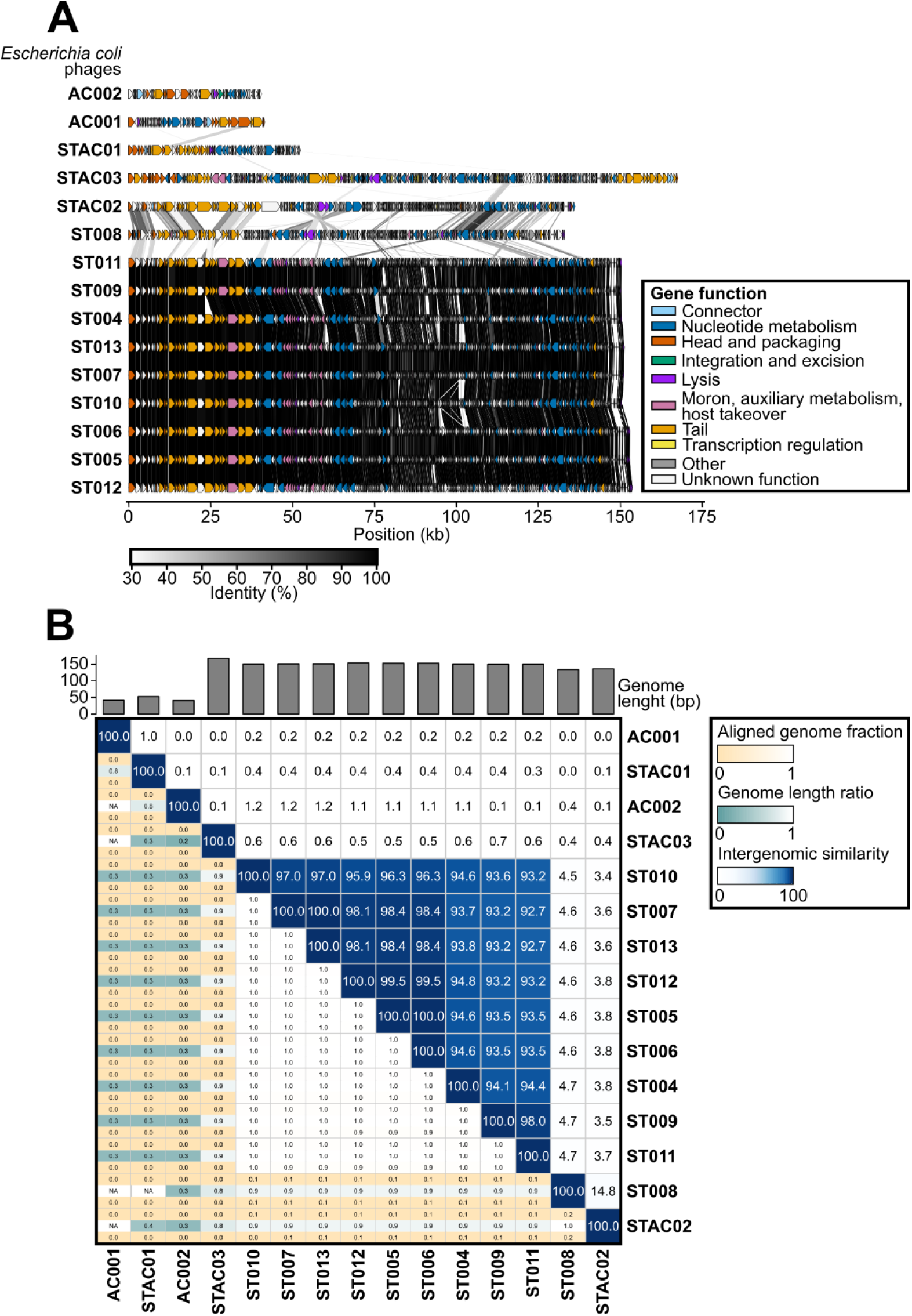
Genomic characterization and comparison of the isolates *E. coli* phages. (A) GenALLYgn (Version 1.0) gene cluster comparison of the isolated phages. Arrows represent genes; each color represents a gene cluster identified by Ally-tool; identity between genes is traced by a gray-to-black gradient from 30 to 100%. (B) Intergenomic heatmap generated by VIRIDIC (BLASTn). Genome length (bp) is shown as grey bars; aliened genome fraction (yellow gradient), genome length radio (green gradient), and intergenomic similarity (blue gradient) are also shown.

The closest relatives of the nine phages belonging to the genus *Phapecoctavirus*, as well as those classified within the genera *Uetakevirus*, *Avunavirus*, *Vequintavirus*, and *Tequatrovirus*, include *Escherichia coli* phages such as ESCO13, PNJ1809-36, vB_EcoM-Ro121|w, vB_EcoP_ZX5, vB_EcoM_TH18, nomo, and vB-Eco-KMB26, respectively. In contrast, phages assigned to the genera *Aghbyvirus* and *Rosemountvirus* showed closer relationship to phages infecting other hosts, including *Serratia* and *Salmonella*, such as vB_SmaP-Kaonashi and emhyr (Table 2).

To further investigate the relationships among the 15 lytic phages, a protein-sharing network analysis was performed. The network was constructed using vConTACT2, based on clusters of shared proteins between our phage genomes and reference phages from the Prokaryotic Viral RefSeq v211 database (Fig. 3A). Phages ΦST005, ΦST006, ΦST012, ΦST011, ΦST013, ΦST007, ΦST009, ΦST010, and ΦST004 formed their own cluster (cluster 57). Phages ΦSTAC02 and ΦST008 clustered closely together in a distinct cluster (cluster 27), which was embedded within the same network neighborhood as cluster 57. Phage ΦSTAC03 was assigned to a separate cluster (cluster 0), which was connected to clusters 57 and 27 through a single reference phage from the database. In contrast, ΦSTAC01 (cluster 205), ΦAC002 (cluster 50), and ΦAC001 (cluster 93) were grouped in very distant clusters with not direct connections to any clusters of the other phages. When comparation were performed among our phages, the nine members of the genus *Phapecoctavirus* formed a distinct cluster, sharing several protein connections with ΦSTAC02 and ΦST008. In contrast, the remaining phages did not share any detectable connection either with this group or among themselves, indicating that ΦSTAC01, ΦAC002, and ΦAC001 are genetically distinct from the other phages (Fig. 3B).

**Figure 3.**
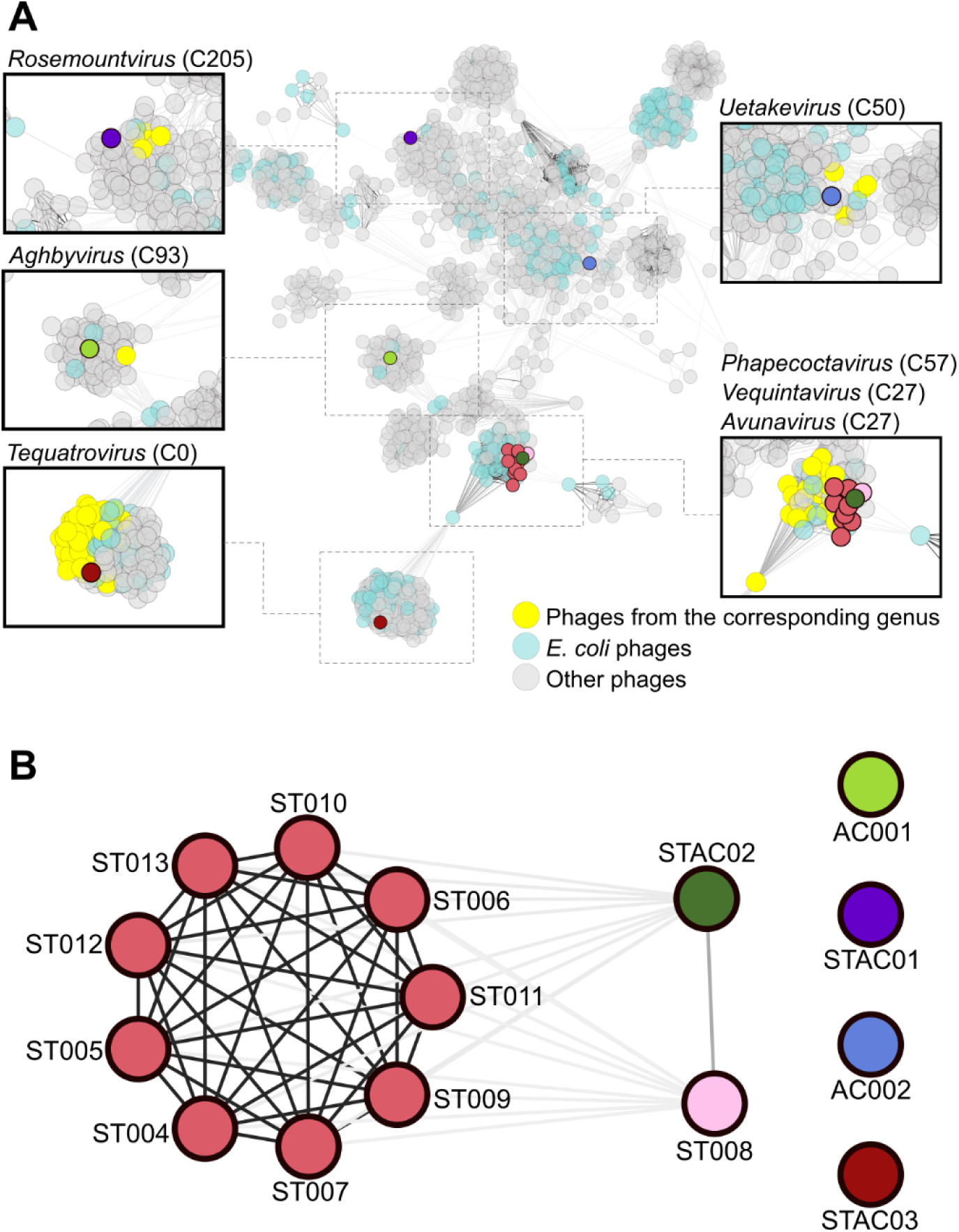
Phages protein-sharing network. (A) vContact2 generated clusters shows that phages are distributed within distinct, genetically distant clusters. Nodes in different colors except yellow, light blue, or grey, represent the isolated phages from this study; nodes in yellow represent phages from specific genus; nodes in light blue represent *E. coli* phages from the Prokaryotic Vira lRefSeq211 database; nodes in grew represent phase from other species excluding *E. coli*. The magnification of the different clusters highlights the genus to which the phage is closest. Number aside the genus name corresponds to the cluster number given by the database. (B) *E. coli* phages described in this study in relation with each other. Phages that cluster together shared the same node color. Edges intensity represents the significance score based on the numbers of proteins shared by the phages.

### Host range analysis

A panel of 16 *E. coli* strains isolated from UTI patients were used to determine the host range of the 15 lytic phages. Host range was assessed by calculating the efficiency of plating (EOP) using the double agar overlay method. The cumulative EOP per strain (CES) and per phage (CEP) were then calculated (Fig. 4). CES values indicated that three strains (1555, 1556, and 1564) were insensitive to all tested phages, whereas strain 1274 was susceptible to only two phages. In contrast, several strains exhibited high cumulative EOP values (e.g., 1554, 1560, and 1566), indicating susceptibility to most or all phages. From the phage perspective, CEP values revealed that ΦAC002 and ΦAC001 infected only two and four strains, respectively, including their original host stain. However, most phages infected between six to twelve strains, with ΦST006, ΦST007, ΦST012, ΦST008, and ΦSTAC03 showing the highest cumulative EOP values. In some cases, CEP values were not proportional to the number of infected strains, suggesting differences in infection efficiency. Specifically, some phages exhibited strong lytic activity in a limited number of strains, whereas others infected a broad range of strains but with lower efficiency. Cocktails design and testing

**Figure 4.**
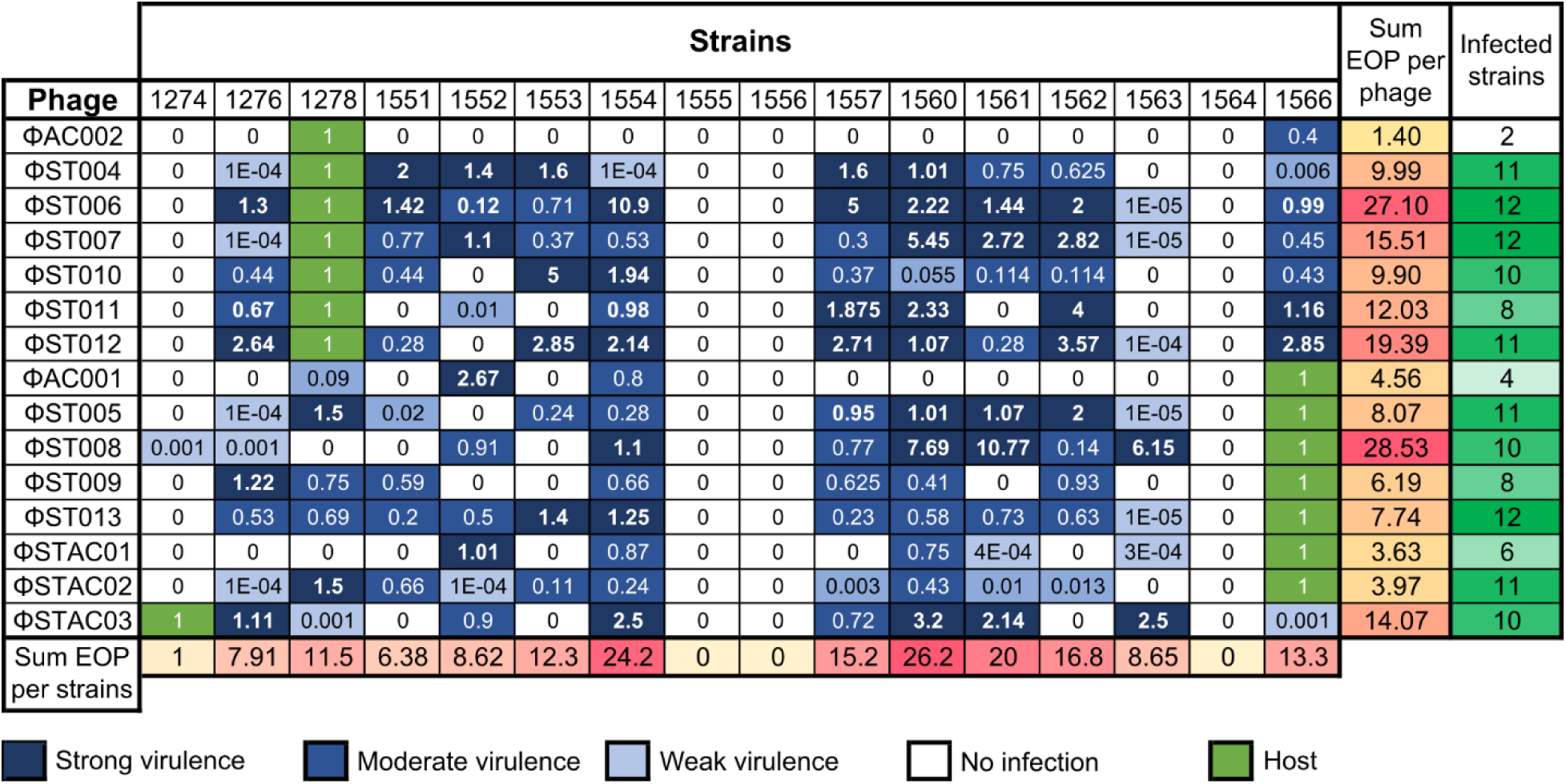
Phage host range and efficiency of plating (EOP). Heatmap showing host range assay for the fifteen isolated phages against eighteen UTI isolated *E. coli* strains at MOI=1. EOP values were classified as strong virulence (EOP > 1 – dark blue); moderate virulence (0.1 ≤ EOP ≤ 1 - blue); weak virulence (0 < EOP < 0.1 – light blue); no infection (EOP = 0 - blank); or propagation host strains (EOP = 1 – green). The cumulative EOP per strain (CES), cumulative EOP per phage (CEP), number of infected strains per phage, and number of infecting phages per strain are shown.

In many clinical settings, bacterial genomic information is not readily available when a phage cocktail must be selected for therapeutic use. Moreover, the identification and annotation of receptor-binding proteins (RBPs) remain challenging due to their high sequence diversity and the large proportion of hypothetical proteins commonly found in phage genomes, some of which may correspond to uncharacterized RBPs. Therefore, to rationally classify phage combinations using a phenotype-based approach, we established a framework based on two key parameters derived from EOP data: host-range overlap and cumulative infection efficiency. Host-range overlap was defined as the proportion of shared susceptible strains (>0.001) between phages within a given combination relative to the total number of strains infected by at least one of the phages (Intersection/Union – Inter/Union). Infection effectiveness (ΣCEP) was estimated using the cumulative EOP values (sum of individual EOP values across all tested strains), which served as a proxy for overall lytic activity. Based on these metrics, we generated sets of phage combinations with distinct profiles of host-range overlap and infection effectiveness. Redundant combinations were intentionally selected to include phages with highly overlapping host range, allowing us to assess whether increased lytic activity could compensate for limited expansion in host coverage. In parallel, complementary combinations were designed by paring phages with partially or minimal overlapping host range, with the aim of increasing the overall breadth of susceptible strains. This strategy allowed us to experimentally test whether functional diversity (low overlap) or lytic strength (high EOP) plays a dominant role in determining cocktails performance (Supp. Table 3).

Using this experimental design, a total of 18 cocktails (C1-C18) were tested in eight *E. coli* isolates from UTI patients, including the 2 host strains (1274 and 1278). Individual phages were also included as controls in the experimental setup (Fig. 5). 24-hour time-killing assays were performed in LB supplemented with MC buffer (LB+MC) to evaluate the lytic activity of each combination. The individual centroid index (CI) by phage or cocktail against each strain was determined, and the cumulative centroid index (CCI) was calculated to assess the overall effect of each phage or cocktail across all the strains. CI values were determined by comparing the growth curves obtained under each experimental condition with their corresponding negative controls with no phages. The number of covered strains (Covered) was determined using a CI threshold of ≥0.3 (Fig. 6). CCI values from the three redundant combinations (C1, C3, and C4) showed additive improvement compared to the single phage components, without any increase in the number of covered strains. In contrast, most complementary combinations exhibited higher CCI values than their individual phages, while maintaining the coverage provided by the most effective single phages. However, some combinations displayed an antagonist behavior, with decreases in both CCI and number of covered strains (C2, C7, C8, and C14), whereas others showed increased CCI values but reduced strain coverage (C9). The most effective combinations were C15 and C18, which demonstrated the highest additive improvement and the greatest CCI values. These results highlight the importance of both complementary and infection efficiency in determining the effectiveness of phages cocktails.

**Figure 5.**
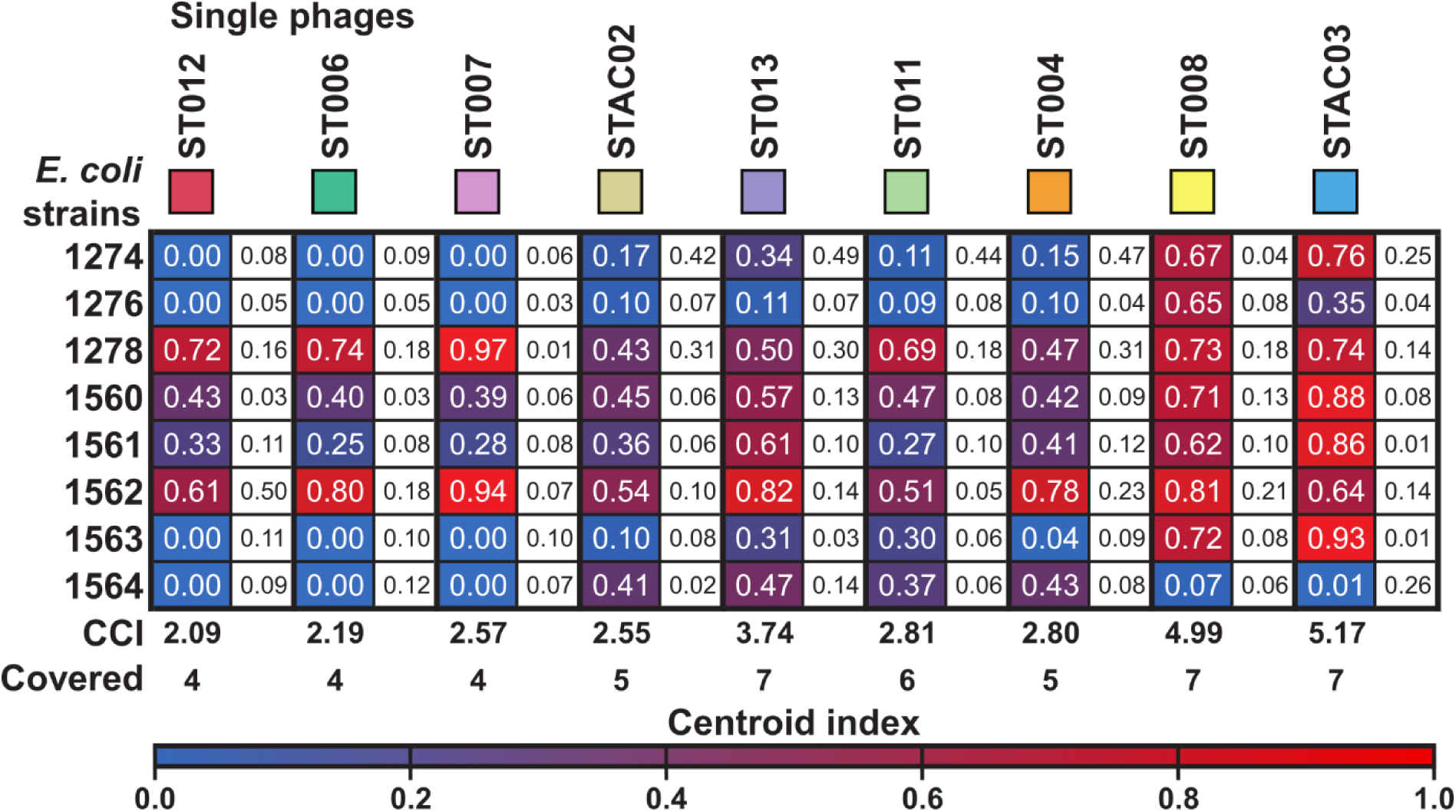
Centroid index of individual phages. Nine individual phages were tested against a panel of eight UTI isolated *E. coli* strains at MOI=1. Heatmap shows the centroid index (CI) values calculated based on 24-hours time-killing curves. CI scale from 1 to 0 is shown in a red-to-blue color gradient. Values were recorded from three independent experiments. Standard deviation (SD) values are shown into the white squares. Cumulative centroid index (CCI) and covered strains (Covered) were calculated.

**Figure 6.**
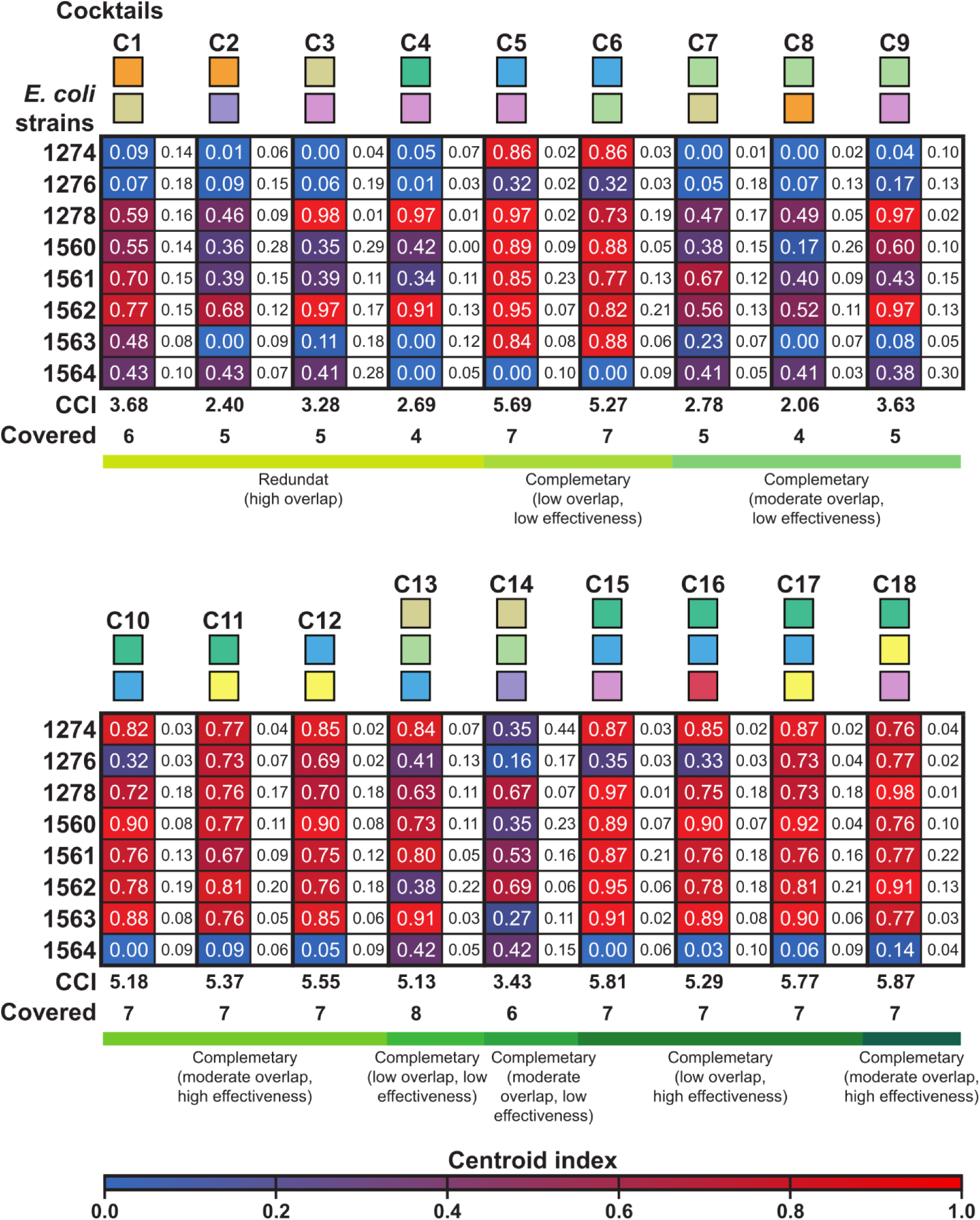
Centroid index of phages cocktails. Twelve dual cocktails, and six triple cocktails were tested against a panel of eight UTI isolated *E. coli* strains at MOI=1. Cocktail classifications are based on Supplementary Table 3 (green gradient). Heatmap shows the centroid index (CI) values calculated based on 24-hours time-killing curves. CI scale from 1 to 0 is shown in a red-to-blue color gradient. Values were recorded from three independent experiments. Standard deviation (SD) values are shown into the white squares. Cumulative centroid index (CCI) and covered strains (Covered) were calculated.

Other aspect that needs to be considered during phage cocktail design is that the bacterial mechanism of phage resistance may involve the downregulation of phage receptor or the activation of anti-phage defense systems, depending on the environment conditions in which the bacteria are naturally growing. Therefore, to better evaluate cocktail performance under conditions that more closely mimic the physiological pathological environment, we tested the two most effective combinations in artificial urine against the eight previously evaluated UTI isolated *E. coli* strains. Time-kill kinetic assays were performed using combinations C15 and C18 in artificial (AU) urine over a 24-hour period. Given that AU is a nutrient-limited medium, heat-inactivated cocktails were included as negative controls to determine the potential contribution of residual nutrients transferred from the phage suspensions. In addition, multiplicities of infection (MOI) of 1.0 and 0.1 were used to determine if the volume of phage lysate added to the cultures could influence the nutrient content into the AU. Experiment was also conducted in LB-LS plus MC buffer by using a MOI of 0.1 as control. The centroid index (CI) and cumulative centroid index (CCI) were subsequently calculated to quantify phage activity. CI values were determined by comparing the bacterial growth curves obtained under each experimental condition with the corresponding negative controls containing heat-inactivated phages (Fig. 7). Regardless of whether an MOI of 1.0 or 0.1 was used, both combinations exhibited markedly lower CCI values in AU than in LB-LS+MC. However, the number of covered strains decreased just in the treatment with cocktail C18. Interestingly, reducing the MOI from 1.0 to 0.1 in LB-LS+MC increased both the CCI values and the number of covered strains for both cocktails.

**Figure 7.**
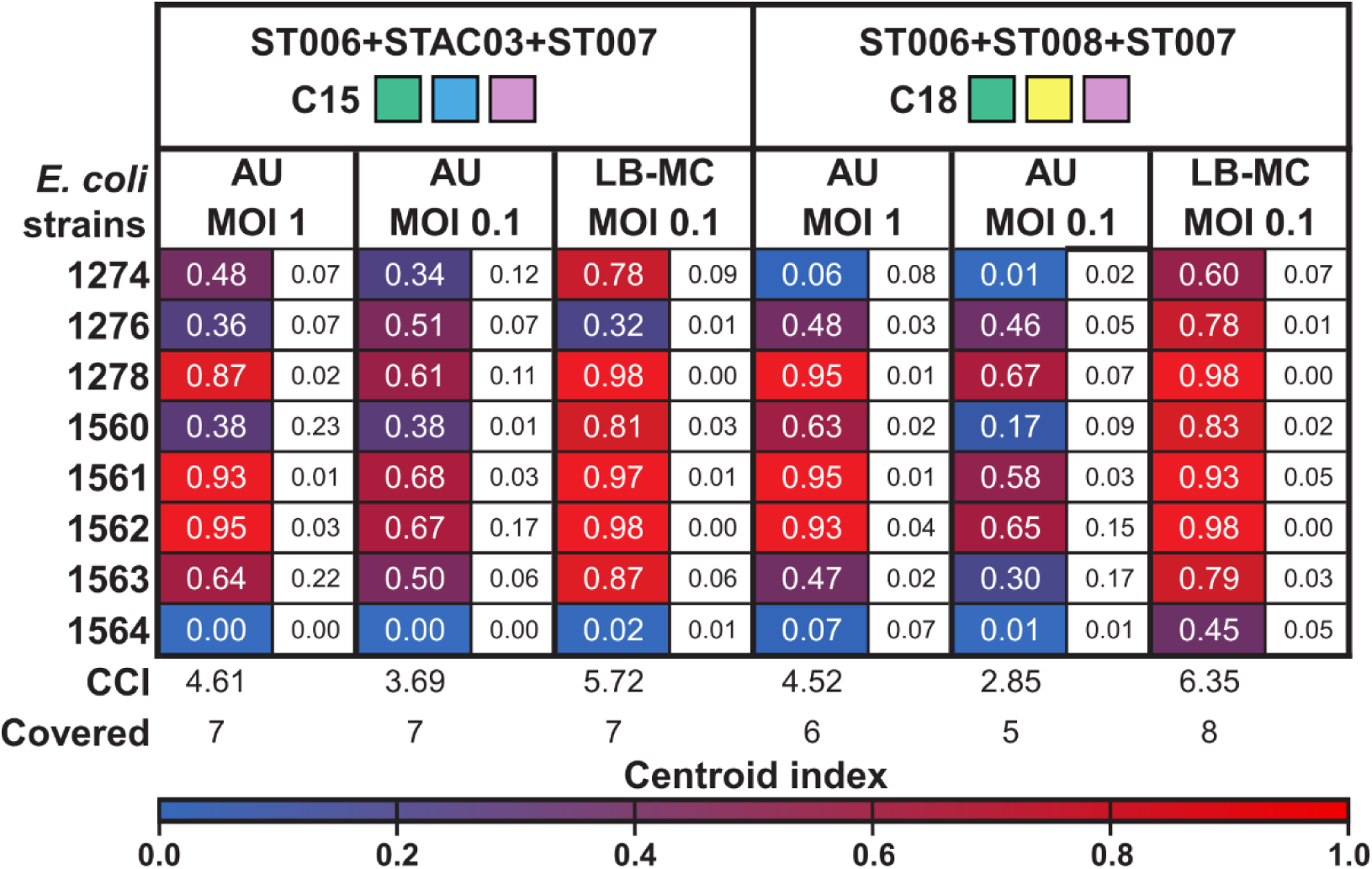
Testing of the most effective cocktails in artificial urine (AU). Cocktails C15 and C18 were tested against a panel of eight UTI *E. coli* isolates in AU at MOI=1.0 and MOI=0.1. Experiment was also set in LB plus MC buffer with a MOI=0.1 as control. Heatmap shows the centroid index (CI) values calculated based on 24-hours time-killing curves. Standard deviation (SD) values are shown into the white squares. Centroid index (CI) was calculated by considering the negative control with the heat-inactive cocktails. Cumulative centroid index (CCI) and covered strains (Covered) were calculated.

## Discussion

One of the major challenges in phage therapy is the selection of effective phages or phage cocktails when limited information about the infecting bacterial strain is available. Although genomic characterization has become increasingly important for understanding phage-host interactions, complete bacterial genome information is rarely available when therapeutic decisions must be made in clinical settings. Conventional diagnostic methods often provide limited genomic resolution (Arbefeville et al., 2024), whereas whole-genome sequencing, despite being considered the gold standard for bacterial characterization, remains relatively expensive, requires specialized bioinformatic infrastructure, and may not provide information rapidly enough for immediate clinical decision-making (Mazhari et al., 2025).

The prediction of phage host range is further complicated by the biology of receptor-binding proteins (RBPs). These highly diverse proteins are frequently subject to recombination and horizontal gene transfer, resulting in receptor specificities that remain poorly characterized for many phages (Pas et al., 2023). In addition, bacterial surface receptors, including lipopolysaccharides, outer membrane proteins, pili, flagella, and extracellular polysaccharides, exhibit substantial diversity among strains (Bertozzi Silva et al., 2016; Moriniere et al., 2026). Consequently, predicting phage susceptibility solely from genomic information remains challenging.

These limitations highlight the need for alternative strategies for phage selection that do not rely exclusively on prior knowledge of bacterial genomic context. Our results suggest that phenotype-based approaches may represent a practical alternative for phage selection in these situations. By integrating host-range overlap and cumulative infection effectiveness, we identified complementary phage combinations capable of maximizing bacterial coverage while minimizing functional redundancy despite the absence of prior knowledge of bacterial receptors or phage RBP specificity. Such strategy may be particularly valuable in clinical scenarios where rapid therapeutic decisions are required and detailed genomic information is unavailable.

Our EOP-based framework enabled the rational design of phage combinations by host-range overlap and infection effectiveness. The results reveled that the most effective cocktails were not necessarily those constituted of the phages with the broadest host ranges or the highest individual EOP (for example ST008 or STAC03). Instead, optimal performance was achieved by combining highly effective phages with broad host range and phages exhibiting high infection efficiency against a more limited set of strains (for example ST006 and ST007). In few cases, the incorporation of a phage with a narrow host-range (specialist) produced an additive effect by enhancing cocktail effectiveness in strains that were only moderately susceptible to the broader-spectrum phages (generalist). This effect was evident in dual-phage cocktails (cocktails C5 and C6) and became even more pronounced in triple-phage combinations (cocktails C15, C16, and C18). Notably, the incorporation of multiple specialist phages into an otherwise broadly active cocktails further increased overall effectiveness, as observed in the case of cocktail 18 (C18). These findings indicate that lytic effectiveness and host-range diversity are independent properties that should be considered separately during phage cocktail design.

A broad host-range alone does not guarantee optimal performance, and highly lytic phages may contribute little if they target the same set of strains. Instead, the most effective cocktails resulted from combining high infection efficiency with functional complementarity, thereby maximizing bacterial coverage while minimizing redundancy. These results suggest that optimal phage cocktails may benefit from combining generalist phages that provide broad coverage with specialist phages that exhibit high infectivity against subsets of otherwise poorly susceptible strains (for example cocktails 15 and 18).

The performance of the redundant cocktails yielded several interesting observations. In most cases, combining phages with highly overlapping host ranges did not result in substantial improvements in infection effectiveness or host coverage (cocktails C1-C4 and C7-C9). Instead, these cocktails either displayed antagonistic effects, resulting in lower effectiveness than the individual phages, or performed similarly to the most effective phage in the combination. These findings indicate that increasing the number of functionally redundant phages does not necessarily improve cocktail performance. One possible explanation is that phages with highly overlapping host ranges may compete for the same share common receptors, thereby limiting the benefits of combining them. Although the mechanisms underlying these interactions were not investigated in this study, the results suggest that redundancy can reduce the functional contribution of individual phages within a cocktail, something that is well described in the literature. These observations have important implications for phage therapy. Larger cocktails may increase production costs, quality-control requirements, and regulatory complexity without providing proportional gains in effectiveness or bacterial coverage. Consequently, cocktail design should prioritize functional complementarity rather than simply maximizing the number of phages included.

An additional consideration is the degree of specialization required to effectively target pathogenic bacteria while minimizing impacts on the surrounding microbiota. Cocktails composed exclusively of broad-host-range phages may increase bacterial coverage but could also affect non-target commensal *E. coli* strains that share common receptors. In this context, the phenotype-based framework proposed here may provide a practical strategy for identifying combinations that maximize effectiveness while avoiding unnecessary expansion of host range.

An additional consideration that should be taken into account long before phage cocktail design is the purity of individual phage isolates. Of the 18 phages initially recovered in this study, three were subsequently identified as mixtures of the target lytic phage and contaminating viral elements, including temperate phages likely originating from prophages present in the propagation hosts. Notably, these contaminants were not evident during isolation or plaque purification and were only detected after genome sequencing. This observation highlights an often-overlooked limitation of phage isolation workflows, particularly when downstream applications rely exclusively on phenotypic characterization.

Prophages can be induced under a wide range of conditions, including environmental stress, antibiotic exposure, and infection by other phages (Goerke et al., 2006; Howard-Varona et al., 2017; Jérôme, 2020). Consequently, induced temperate phages may involuntarily contaminate the isolation and propagation of lytic phages, potentially influencing host-range determination, EOP measurements, and cocktail performance. This issue is particularly relevant given that several phage therapy studies report biological characterization of therapeutic phages without providing genome sequences or detailed assessments of isolate purity (Jault et al., 2019; Law et al., 2019; Leitner et al., 2021; Rostkowska et al., 2021; Wu et al., 2021; Zaldastanishvili et al., 2021). Under such circumstances, the observed antibacterial activity could originate from the target phage, an induced prophage, or the combined activity of multiple viral elements. Our findings therefore support genome sequencing not only as a tool for taxonomic classification and genome characterization, but also as an essential quality-control step for confirming phage identity and purity prior to therapeutic development and cocktail design.

One of the primary challenges in phages therapy is the ongoing cycle of co-evolution between bacterial hosts and phages. Mechanisms of phage resistance include down-regulation of phage receptors, action of restriction modification enzymes, CRISPR-Cas adaptive immunity, and spontaneous mutations (Labrie et al., 2010). Importantly, the expression and regulation of these mechanisms are often influenced by environmental conditions. For example, the porins OmpC and OmpF, which serve as receptors for several E. coli phages, are differentially expressed according to medium osmolarity, with OmpC predominating under high-osmolarity conditions and OmpF under low-osmolarity conditions (Hantke, 2020; Yoshida et al., 2006). Lipopolysaccharide (LPS), another common phage receptor, can vary depending on nutrient availability, including glucose, phosphorus, or magnesium availability (Javed et al., 2023).

Most phage characterization studies are performed in Luria-Bertani (LB) medium, a nutrient-rich environment optimized for bacterial and phage propagation, especially in *E. coli* (Lisac & Podgornik, 2025). In contrast, urine composition change depending on many factors, such as age, gender, race, and food intake, and even the time of collecting the sample from the same person can affect the composition (Ferrão et al., 2024). All these constantly changing aspects in human urine could altered the expression of bacterial surface structures involved in phage adsorption, thereby affecting phage infectivity and resulting in host-range patterns that differ from those observed in standard laboratory media.

Consistent with this hypothesis, both combinations C15 and C18 showed substantially reduced activity in AU compared with LB-LS+MC, as evidenced by lower CCI values and reduced strain coverage. Furthermore, heat-inactivated cocktails used as controls had little effect on bacterial growth, suggesting that the observed decrease in activity was not simply due to nutrient limitation or nutrient carryover from the phage lysates. Together, these observations indicate that factors intrinsic to the urinary environment may induces a stronger influence on phage efficacy.

These results support the view that bacterial physiology, and consequently receptor availability, may differ substantially between laboratory media and physiologically natural environments. Therefore, determining phage activity exclusively in rich culture media may overestimate therapeutic performance under urinary tract conditions. Therefore, to better investigate phage-host interactions in a context that resembles the physiological environment, we established and evaluated a standardized artificial urine (AU) formulation based on that reported by Rimbi *et al*. in 2024 (Rimbi et al., 2024) . Such approach may provide a more representative platform for phage screening, cocktail optimization, and the prediction of therapeutic efficacy against uropathogenic *E. coli*. Additionally, future studies combining receptor mapping and transcriptomic approaches will be necessary to determine the mechanistic basis underlying the variations observed between laboratory and urine-based media.

Several limitations of this study should be acknowledged. First, although our results suggest that environmental conditions influence phage activity through changes in bacterial physiology, the specific receptors involved in phage adsorption were not experimentally identified, and the proposed framework does not explicitly incorporate receptor usage, RBP diversity, or the molecular basis of host recognition. Second, host-range and cocktail performance were evaluated using a limited collection of UTI-associated *E. coli* isolates, and validation in larger and more diverse strain collections will be necessary to determine the broader applicability of this approach. Third, although artificial and natural urine provide physiologically relevant environments, the experiments remained in vitro and do not fully reproduce the complexity of urinary tract infections. Finally, the detection of prophage-derived contaminants in several initially isolated phages highlights the importance of genomic validation during phage discovery and cocktail development. Despite these limitations, our results demonstrate that host-range overlap and cumulative infection effectiveness provide a practical phenotype-based framework for identifying complementary phage combinations, particularly in situations where detailed bacterial genomic information is unavailable. This approach successfully identified highly effective cocktails under physiologically relevant conditions and may contribute to the development of more rational strategies for phage therapy against urinary tract infections.

## Methodology

### Bacterial strains and growth conditions

All *Escherichia coli* strains used in this study are listed in Supplementary Table 1. *E. coli* clinical strains were collected from UTI patients from the Centre Hospitalier Universitaire de Sherbrooke (CHUS), located at Sherbrook, Quebec, Canada. Mice *E. coli* strain 1566 was isolated from feces from a heathy mouse breaded at IRCUS (Institut de recherche sur le cancer de l’Université de Sherbrooke). Luria-Bertani Low Salt (LB-LS) (Sigma-Aldrich, Cat. No. L3397-1KG) was used for all the experiments excluding those performed in Artificial Urine. Bacteria were streaked onto LBLS-agar plates and grown overnight at 37°C. One single colony was picked up and inoculated in LB-LS broth and incubated overnight at 37°C by shaking at 220 rpm.

### Phage isolation and purification

Wastewater samples were collected from different sources, including rivers, streams, dams, vegetable fields, swamps, wastewater treatment plants, and the Jeffrey-Gingras Park, all located in Sherbrooke, Quebec, Canada. Samples were centrifuged at 3220 g and pre-filtered by passing them through a conventional paper filter to remove unwanted debris. Samples were re-filtered through a 0.22-micron filter in order to remove all remaining bacteria. Filtered samples were stored at 4°C until use.

To isolate and purify environmental phages, 9 ml of filtered sample were mixed with 1 ml of LB 10X, followed by adding 100 ul of overnight culture of the sensitive bacteria plus MC buffer (MgCl_2_ and CaCl_2_) to a final concentration of 10 mM. The mixtures were incubated overnight at 37°C by shaking at 220 rpm. After incubation, the mixtures were centrifuged at 3220 g for 20 minutes at 4°C. Supernatants were then filtered through a 0.22-micron filter to remove residual bacteria.

Infecting phages were detected by double-layer agar method. For this, 100 ul of filtered supernatants and 100 ul of overnight culture of the sensitive bacteria were mixed with 3 ml of soft agar (0.3%) plus MC buffer (10 mM final). The mixtures were poured onto 1% LB agar plates and incubated overnight at 37°C. The presence of phages was confirmed by the formation of clear lysis plaques on the bacterial lawn. Putative single plaques were picked up and re-suspended in phage buffer (50 mM Tris-HCl pH 7.5; 100 mM NaCl; 8 mM MgSO_4_), then incubate at room temperature for 3 h to let the phages diffuse to the buffer. This cycle of steps was repeated at least three times until plaques morphology was homogeneous for each phage.

### Phage amplification and titration

For phages amplification, the same process mentioned before for phage enrichment from environmental samples was done with the only difference of using LB-LS 1X. The rest of the process is the same as described earlier. Phage titration was obtained by spot test and double-layer agar method described before. Ten-fold serial dilutions were performed in LB-LS plus MC buffer, followed by spotting 10 ul of dilutions onto the soft agar. Plates were left to dry and incubated overnight at 37°C. Phage titers were verified before other experiments. Stocks were stored at 4°C.

### Phage DNA genomic extraction

Phage were concentrated by polyethylene glycol (PEG) precipitation at first. Precipitation was performed by mixing 10% wt/vol of PEG and 2 mL of NaCl 5M with 10 ml of phage lysate. Samples were incubated at 4°C overnight. After incubation, mixtures were centrifuged at 20,817 g for 30 minutes to pellet the phages. Pellets were resuspended in 400 ul of phage buffer 1X and subjected to treatment with DNase and RNase (2,000 units) for 30 minutes at 37°C to remove any contamination with bacterial DNA. Treatment with nucleases were repeated twice. DNA extraction was performed with the Monarch Spin Genomic DNA Extraction kit (New England Biolabs, Cat. No. T3010S) by following manufacturer instructions. DNA concentration was quantified by DS-11 series spectrophotometer (DeNovix). DNA integrity, purity, and size were visualized by agarose gels (0.8%).

### Bioinformatic analysis of the lytic phages

Whole genome sequencing of the phages was done by using a PromethION 2 solo sequencer (Oxford nanopore technology). Library preparation, sequencing, and genome assemblies was performed by the RNomic platform of Université de Sherbrooke (https://rnomics.med.usherbrooke.ca/).

Genomes annotation was conducted by Pharokka (version 1.8.2) (Bouras et al., 2023) with its database (version 1.8.0). Phold (version 0.2.0) and Empathi (Bouras et al., 2026) were also used to enhance genome annotations (exact parameters can be found at https://github.com/allychamp/phage-genome-analysis-pipeline). Protein-sharing network was constructed by vConTACT2 (version 0.11.3) (Bolduc et al., 2017; Jang et al., 2019) using the Prokaryotic Viral RefSeq 211-Merged database and visualized in Cytoscape 3.10.4 (Shannon et al., 2003). Intergenomic heatmap was generated by VIRIDIC (BALSTn) (Moraru et al., 2020). GenALLYgn (Version 1.0.0) code is available at GitHub https://github.com/allychamp/GenALLYgn.

### Host-range spectrum and efficiency of plating (EOP)

Host range was determined using the standard spot assay described above. Phages were tested against a panel of UTI-associated E. coli isolates (Supplementary Table 1). Efficiency of plating (EOP) was calculated as the ratio between the phage titer obtained on the target strain and the titer obtained on the propagation host strain. EOP values were classified as strong virulence (EOP > 1), moderate virulence (0.1 ≤ EOP ≤ 1), weak virulence (0 < EOP < 0.1), or no infection (EOP = 0), corresponding to the absence of visible plaques. The cumulative EOP per strain (CES), cumulative EOP per phage (CEP), number of infected strains per phage, and number of infecting phages per strain were subsequently calculated.

### Cocktail design

Phages with the highest cumulative EOP per phage (CEP) and/or larger host range (covered strains) were chosen for phage cocktails. Dual and triple cocktails were designed by Fractional Factorial Desing (FFD) to systematically evaluate phage combinations while reducing the number of experimental conditions required for a full factorial design. Phage candidates were treated as binary factors according to their presence or absence in each cocktail. Selected cocktails were tested against the target bacteria in time-killing assays performed in triplicates.

### Time-killing assays

24-hour time-killing assays were conducted in 96-well plates with a final volume of 200 ul. 20 ul of bacteria cultures at an OD_600_ of 1 (∼1x10^9^ CFU/ml) were diluted in 130 ul of LB-LS plus MC buffer to have ∼1x10^7^ CFU/ml. At the same time, ∼1x10^7^ PFU/ml of individual phages or combinations were added to the mix to obtain a multiplicity of infection (MOI) of 1.0. Final volume was adjusted to 200 ul with LB plus MC buffer. Bacterial growth was monitored by measuring the optical density at a 600 nm (OD_600_) by a BioTek-Epoch2 plate reader (Agilent). Plates were incubated at 37°C and measurements were recorded every 5 minutes for 24 hours. Before each measurement, reactions were mixed by orbital shaking for 30 seconds. Results were compiled with the Gen5 software (Agilent). Experiments were conducted in triplicate in three independent experiments. Centroid index (CI) (Hosseini et al., 2024), cumulative centroid index (CCI), and covered strains (host-range) of the phages alone or in combinations were calculated.

### Validation of cocktails in artificial urine (AU)

Artificial urine was adapted from publish formulations with small changes (Rimbi et al., 2024; Sarigul et al., 2019). Components, stock concentrations, and store conditions are shown in Supplementary table 4. All stocks were prepared in distilled deionized water and filtered sterilized by 0.22-micron filter except for peptone that was sterilized by autoclave. Unlike the published formulations, peptone was used at 0.03%, thiamine hydrochloride at 1 mg/L, and dextrose anhydrous at 0.1886 mmol/L. Urea was weighted and added during the mixing process. Artificial urine was stored up to one week at 4°C until used.

24-hour time-killing assays were conducted as mentioned before with few changes. Experiments were performed in AU or AU plus MC buffer. From each strains tested, ∼1x10^9^ CFU/ml were wash with saline (NaCl 0.9%) by two cycles of centrifugation (10 min at 8,000 rpm at room temp) and resuspension. After that, bacteria were resuspended in AU. 20 ul of bacteria cultures at were diluted in 130 ul of AU to have at final ∼1x10^7^ CFU/ml. Individual phages or in combination were used at ∼1x10^6^ CFU/ml (MOI 0.1). A MOI of 1.0 and 0.1 was used to compare the effect of transferring unwanted nutrients from the phage lysates into the AU. Negative controls without phages and heat-inactive phages were used. Heat-inactivation was performed at 90°C for 60 minutes. Final volumes were adjusted to 200 ul. Growth monitoring was recorded as in common time-killing assays previously mentioned. Experiments were conducted in triplicate in three independent experiments. Centroid index (CI) was calculated by considering the curves from the negative control with heat-inactive cocktails, as well as the testing experiment. Cumulative centroid index (CCI) and covered strains (host-range) of the phages alone or in combinations were also calculated.

## Statements

### Data availability statement

GenBank accession numbers are listed in Table 2.

### Author contributions

Paul Ugalde Silva: Formal analysis, Investigation, Methodology, Validation, Data curation, Writing – original draft, Writing – review and editing. Ally Champoux: Formal analysis, Investigation, Software, Data curation, Writing – original draft. Annie Chénard: Formal analysis, Investigation, Methodology, Validation. Serena Tuytschaevers: Formal analysis, Investigation, Validation. Élodie Jolicoeur: Methodology, Validation. Nathalie Rivard: Resources, Writing – review and editing. Louis-Charles Fortier: Conceptualization, Funding acquisition, Project administration, Resources, Supervision, Writing – original draft, Writing – review and editing.

## Acknowledgments

We thank the RNomics Platform of the U. Sherbrooke for genome sequencing. A. Champoux and E. Jolicoeur are both recipients of a Canada Graduate Research Scholarship (CGRS M) from the Natural Sciences and Engineering Council of Canada (NSERC), and a research scholarship from the Fonds de Recherche du Québec – Secteur Nature et Technologies; A. Chénard was the recipient of a “Bourse d’Excellence en Recherche” from the U. Sherbrooke; S. T. was the recipient of a Fulbright Canada-Mitacs Globalink Research scholarship. This work was supported by a NSERC Discovery Grant (RGPIN-2020-05776, to L.C. Fortier), Canadian Institutes of Health Research grants (202309ARB and 202503PJT to L.C. Fortier, and 202303-PJT to N. Rivard); and a Canada Foundation for Innovation Grant John R. Evans Leaders Fund (to É. Massé, U. Sherbrooke). (L.C. Fortier is a member of the Centre de Recherche du CHUS.

## Notes

### Competing Interest Statement

The authors have declared no competing interest.

