## Supplementary Table 1 for "Genomic and phenotype-based framework for rational design of phage cocktails against *Escherichia coli* isolates from urinary tract infections"

**Supplementary Table 1. List of bacteria used in this study.**

| ***Escherichia coli* strain** | **Source** |
|---|---|
| 1566 | Commensal derived from laboratory mouse |
| 1274 | Clinical isolate from a patient with UTI |
| 1276 | Clinical isolate from a patient with UTI |
| 1278 | Clinical isolate from a patient with UTI |
| 1551 | Clinical isolate from a patient with UTI |
| 1552 | Clinical isolate from a patient with UTI |
| 1553 | Clinical isolate from a patient with UTI |
| 1554 | Clinical isolate from a patient with UTI |
| 1555 | Clinical isolate from a patient with UTI |
| 1556 | Clinical isolate from a patient with UTI |
| 1557 | Clinical isolate from a patient with UTI |
| 1560 | Clinical isolate from a patient with UTI |
| 1561 | Clinical isolate from a patient with UTI |
| 1562 | Clinical isolate from a patient with UTI |
| 1563 | Clinical isolate from a patient with UTI |
| 1564 | Clinical isolate from a patient with UTI |
