## Supplementary Table 2 for "Genomic and phenotype-based framework for rational design of phage cocktails against *Escherichia coli* isolates from urinary tract infections"

**Supplementary Table 2. Isolate phages considered as contaminated.**

| *Escherichia coli* phage | Genome size (bp) | CDS | Lysogeny-associated genes | Closest relative (INPHARED) | Lowest taxa (genus) | Identity (%) | Coverage (%) | GC content (%) |
|---|---|---|---|---|---|---|---|---|
| ST014 | 136,284 | 254 | 0 | *Escherichia* phage nomo | *Vequintavirus* | 97.9 | 99 | 44 |
| | 151,093 | 307 | 0 | *Escherichia* phage ESCO13 | *Phapecoctavirus* | 98.35 | 95 | 39 |
| | 43,389 | 68 | 2 | Enterobacteria phage mEp460 | *Glaedevirus* | 97.11 | 84 | 51 |
| AC003 | 40,632 | 62 | 2 | *Escherichia* phage vB_EcoP_ZX5 | *Uetakevirus* | 99.14 | 81 | 49 |
| | 97,863 | 154 | 1 | Punavirus P1 | *Punavirus* | 98.9 | 82 | 48 |
| AC004 | 40,615 | 62 | 2 | *Escherichia* phage vB_EcoP_ZX5 | *Uetakevirus* | 98.71 | 82 | 49 |
| | 97,986 | 156 | 1 | Punavirus P1 | *Punavirus* | 98.63 | 75 | 48 |
