## Supplementary Table 3 for "Genomic and phenotype-based framework for rational design of phage cocktails against *Escherichia coli* isolates from urinary tract infections"

**Supplementary Table 3. Phage cocktail design**

| No. cocktail | Combination | Intersection (common strains) | Union (covered strains) | Overlap (Inter/Union) | Effectiveness (ΣCEP) | Classification |
|---|---|---|---|---|---|---|
| C1 | STAC02+ST004 | 8 | 10 | 0.8 | 13.96 | Redundant (high overlap) |
| C2 | ST004+ST013 | 9 | 11 | 0.82 | 17.73 | Redundant (high overlap) |
| C3 | STAC02+ST007 | 9 | 10 | 0.9 | 19.48 | Redundant (high overlap) |
| C4 | ST006+ST007 | 10 | 11 | 0.909 | 42.61 | Redundant (high overlap) |
| C5 | STAC03+ST007 | 5 | 13 | 0.385 | 29.582 | Complementary (low overlap, low effectiveness) |
| C6 | STAC03+ST012 | 5 | 13 | 0.385 | 33.462 | Complementary (low overlap, low effectiveness) |
| C7 | ST011+STAC02 | 6 | 11 | 0.55 | 15.99 | Complementary (moderate overlap, low effectiveness) |
| C8 | ST011+ST004 | 6 | 11 | 0.55 | 22.02 | Complementary (moderate overlap, low effectiveness) |
| C9 | ST011+ST007 | 7 | 11 | 0.64 | 27.54 | Complementary (moderate overlap, low effectiveness) |
| C10 | ST006+STAC03 | 6 | 13 | 0.462 | 41.172 | Complementary (moderate overlap, high effectiveness) |
| C11 | ST006+ST008 | 7 | 12 | 0.583 | 55.632 | Complementary (moderate overlap, high effectiveness) |
| C12 | STAC03+ST008 | 6 | 10 | 0.6 | 42.604 | Complementary (moderate overlap, high effectiveness) |
| C13 | STAC02+ST011+STAC03 | 3 | 13 | 0.23 | 30.06 | Complementary (low overlap, low effectiveness) |
| C14 | STAC02+ST011+ST013 | 6 | 11 | 0.55 | 23.73 | Complementary (moderate overlap, low effectiveness) |
| C15 | ST006+STAC03+ST007 | 5 | 13 | 0.385 | 56.682 | Complementary (low overlap, high effectiveness) |
| C16 | ST006+STAC03+ST012 | 5 | 13 | 0.385 | 60.562 | Complementary (low overlap, high effectiveness) |
| C17 | ST006+STAC03+ST008 | 5 | 13 | 0.385 | 69.704 | Complementary (low overlap, high effectiveness) |
| C18 | ST006+ST008+ST007 | 7 | 12 | 0.583 | 71.142 | Complementary (moderate overlap, high effectiveness) |
