## Supplementary Table 4 for "Genomic and phenotype-based framework for rational design of phage cocktails against *Escherichia coli* isolates from urinary tract infections"

**Supplementary Table 4. Artificial urine formulation based on Rimbi *et al*., 2024 and Sarigul *et al*., 2019.**

| Components | Stock concentration | Molarity (g) | Concentration/L | Storage location | For 100 ml of AU (ul) |
|---|---|---|---|---|---|
| Sodium sulfate | 1M | 142.04 | 11.965 mmol/L | RT | 1196.5 |
| Trisodium citrate | 1M | 258.07 | 2.45 mmol/L | 2-8°C | 245 |
| Creatinine | 1M | 113.12 | 7.791 mmol/L | - 20°C | 779.1 |
| Urea | Weight directly | 60.06 | 249.75 mmol/L | N/A | 1.49 g |
| Potassium chloride | 1M | 74.55 | 30.953 mmol/L | RT | 3095.3 |
| Sodium chloride | 1M | 58.44 | 30.953 mmol/L | RT | 3095.3 |
| Calcium chloride | 1M | 147.01 | 1.663 mmol/L | RT | 166.3 |
| Ammonium chloride | 1M | 53.49 | 23.667 mmol/L | RT | 2366.7 |
| Potassium oxalate monohydrate | 1M | 184.23 | 0.19 mmol/L | RT | 19 |
| Magnesium sulfate heptahydrate | 1M | 246.47 | 4.389 mmol/L | RT | 438.9 |
| Sodium phosphate monobasic dihydrate | 1M | 156.01 | 18.667 mmol/L | RT | 1866.7 |
| Sodium phosphate dibasic dihydrate | 0.5M | 177.99 | 4.667 mmol/L | RT | 933.4 |
| Dextrose (D-Glucose) Anhydrous | 0.1M | 180.16 | 0.1886 mmol/L | RT | 188.6 |
| Uric acid | 0.5M | 168.11 | 0.25 g/L | - 20°C | 297.4 |
| Peptone | 10% | N/A | 0.03% | RT | 250 |
| Thiamine hydrochloride | 1 mg/ml | N/A | 1 mg/L | - 20°C | 100 |

**Note: RT - room temperature; N/A - Not applicable.**
